# Implications of source-sink feedbacks for modelling tree carbon assimilation and growth

**DOI:** 10.1101/2024.09.27.615358

**Authors:** Andrew D. Friend, Yizhao Chen, Annemarie H. Eckes-Shephard, Patrick Fonti, Eva Hellmann, Tim Rademacher, Andrew D. Richardson, Patrick R. Thomas

**Affiliations:** Department of Geography, University of Cambridge, Downing Place, Cambridge, CB2 3EN, UK; Conservation Research Institute, University of Cambridge, Pembroke Street, CB2 3QZ, UK; College of Ecology and Environment, Nanjing Forestry University, Nanjing, China; Department of Physical Geography and Ecosystem Science, Lund University, Sölvegatan 12, S-223 62, Sweden; Research Unit Forest Dynamics, Swiss Federal Institute for Forest Snow and Landscape Research WSL, Zürcherstrasse 111, Birmensdorf, 8903, Switzerland; Sainsbury Laboratory University of Cambridge, CB2 1LR, UK; Institute of Temperate Forest Sciences, University of Quebec in Outaouais, 58 Rue Principale, J0V 1V0, Canada; Center for Ecosystem Science and Society, Northern Arizona University, Arizona, USA; School of Informatics, Computing, and Cyber Systems, Northern Arizona University, Arizona, USA; Department of Pure Mathematics and Mathematical Statistics, University of Cambridge, Wilberforce Road, CB3 0WB, UK

**Keywords:** growth, model, source, sink, feedback, homeostasis, sucrose

## Abstract

Current global models of vegetation dynamics are largely carbon (C) source-driven, with behaviour primarily determined by the environmental responses of photosynthesis. However, real plants operate as integrated wholes, with feedbacks between sources, such as photosynthesis, and sinks, such as growth, resulting in homeostatic concentrations of metabolites such as sugars. A parsimonious approach to implementing this homeostatic coupling of C sources and sinks in a tree growth model is presented, and its implications for the responses of net photosynthesis and growth to environmental factors and tree size assessed. Hill functions describe inhibition of C sources (net photosynthesis) and activation of sinks (structural growth) as sucrose concentration increases. The model is parameterised for a typical tree growing at a site in the Amazonian rainforest and its qualitative behaviour is found to be consistent with observations. A key outcome is that sinks and sources strongly regulate each other. Hence environmental factors that affect potential net photosynthesis, such as atmospheric CO_2_, have greatly reduced effects on growth when homeostatic feedbacks from sucrose concentrations are considered. For example, compared with a C-source-only-driven approach (as in most current global models), the response of tree biomass for a tree currently 300 yr old, to increasing atmospheric CO_2_ projected to the end of this century under a high scenario, is reduced by ca.77%, from +122% to +29%, with net photosynthesis and growth rate responses reduced by a similar amount. Furthermore, in this coupled approach, any direct controls on growth (either environmental or through phenological controls on xylogensis) will influence source activity through the sucrose feedback. For example, a reduction in potential growth through temperature constraints on cell-wall construction increases sucrose concentrations, resulting in a compensating reduction in net photosynthesis. While net photosynthesis controls growth, growth controls net photosynthesis. In addition, we find a strong effect of changing tree allometry on C source-sink relations as the tree grows. Larger trees are less source-limited due to a higher ratio of sapwood area (and hence potential C assimilation rate) to potential growth rate, consistent with the observed decline in growth response to atmospheric CO_2_ as trees age. We suggest that the implications of including C source-sink coupling in models of vegetation dynamics, such as dynamic global vegetation models, are likely to be profound.

## 1 Introduction

Plant growth can be characterised as an increase in cell number, biomass, and/or volume (Hilty et al. 2021). Growth determines the resource acquisition, competitivity, and reproductive output of individuals. Plant growth also plays a major role in the global carbon (C) cycle, estimated to sequester at least 80 Pg[C] yr^*−*1^ in terrestrial biomass annually (Graven et al. 2011), over 7x the estimated total anthropogenic emissions of C in 2023 (Friedlingstein et al. 2023).

Unlike in animals, growth in plants is typically modular and indeterminate (Brukhin and Morozova 2011). Growth requires a supply of resources, and the movement of carbohydrates and other nutrients within plants to support growth is commonly characterised as occurring from ‘sources’ to ‘sinks’ (Mason and Maskell 1928). C is mainly transported as a component of sucrose (Dominguez and Niittylä 2022), with mature leaves being the principal source organs. C sinks include active meristems, which import assimilates for growth and reproduction, as well as processes such as respiration and exudation (Kaitaniemi and Honkanen 1996; White et al. 2016). Storage tissues such as parenchyma can be a C source or sink, depending on the plant-level balance of carbohydrate supply and demand (Furze et al. 2019; MacNeill et al. 2017; Saddhe et al. 2021).

The mechanisms determining the relationships between source and sink activities within plants, and the balance between different sinks, have been researched for many decades (e.g. Wardlaw 1990; Hartmann et al. 2020). Much of this work has been driven by the desire to improve crop production through reducing limitations to growth (e.g. Wardlaw 1990; White et al. 2016; Fernie et al. 2020), but the underlying mechanisms determining source-sink relationships are also highly relevant to the future viability and productivity of plants growing in natural ecosystems in the context of environmental change (e.g. Hartmann et al. 2020; Shestakova et al. 2024), and the dynamics of the global carbon cycle (e.g. Fatichi et al. 2019; Friend et al. 2019). While understanding of carbohydrate metabolism is much more advanced in small herbaceous annuals such as *Arabidopsis* than large woody perennials such as trees (Wingler and Henriques 2022), a key on-going debate in both agronomic and global-change research concerns whether C sources or sinks, especially growth sinks, are ‘in control’ of dry matter production (e.g. White et al. 2016; Körner 2015; Camarero and Andrés 2014; Gessler and Zweifel 2024).

Sucrose performs three key roles at the whole-plant scale: (1) a store of energy and a vehicle for its supply; (2) a metabolic precursor and supplier of building blocks; and (3) regulatory signalling (Fichtner et al. 2021; Saddhe et al. 2021; Choudhary et al. 2022; Göbel and Fichtner 2023). While the process is not fully understood, especially in trees, sucrose is also responsible for producing the hydrostatic gradient that drives phloem transport (Schepper et al. 2013). A major reason that sucrose, rather then glucose, is used by plants for long-distance transport is because it is non-reducing and therefore relatively metabolically inert, protecting critical proteins from non-enzymatic glycosylation (Geiger 2020). Presumably because of its central role in numerous critical processes, complex regulatory pathways maintain homeostasis in sucrose levels (Miret et al. 2024), and are the focus of much current research (e.g. Avidan et al. 2024; Chang and Zhu 2017). ‘Homeostasis’ signifies a self-regulating system in which feedbacks sensitive to its state result in some form of stability of that state relative to a perturbation. We do not infer homeostasis to imply any particular degree of stability. Therefore the system’s state can change in response to varying conditions, even at equilibrium, but feedbacks will result in less change in state than if they were absent. Homeostasis in sucrose concentrations ensures energy and precursor supply rates to cells are compatible with their metabolic capacities and that overall osmotic balance is maintained (Saddhe et al. 2021; Dominguez and Niittylä 2022). While many details of the regulatory networks responsible for sucrose homeostasis remain unknown, feedbacks on source and sink activities clearly play essential roles (e.g. Avidan et al. 2024).

Promotion of C-source activities, such as photosynthesis and remobilisation from storage tissues, occurs when sugars are depleted, whereas promotion of sink activities, such as cellular growth and biosynthesis, occurs when sugars are abundant (Eveland and Jackson 2012; Avidan et al. 2024). Boussingault (1868) is reported to have been the first to postulate that photosynthesis is regulated by its end products (Neales and Incoll 1968; Paul and Foyer 2001). Subsequent work has frequently observed feedback regulation of photosynthesis (Ainsworth and Bush 2011), which has been linked to whole-plant nitrogen status (Paul and Foyer 2001), although it may not occur when it is drought-stress that reduces growth (Gessler and Zweifel 2024), possibly because drought also directly controls photosynthesis (Thompson et al. 2023) and/or may increase demand for assimilates through a stress response (McDowell 2011). Sucrose can be sensed directly, or a signal is expressed through glucose or UDP-glucose products, with these hexoses promoting organ growth and cell proliferation, while sucrose itself promotes differentiation and maturation (Borisjuk et al. 2002; Koch 2004). The relevance of these different regulatory roles in tree growth can be seen in relation to differential spatial distributions of metabolites across developing wood (e.g. Uggla et al. 2001).

Evidence for source-sink feedbacks operating through these mechanisms in mature trees has been found by altering rates of C-supply to the cambial meristem using phloem girdling, compression, and chilling (Rademacher et al. 2021; Rademacher et al. 2022). Rademacher et al. (2021), using phloem girdling and compression treatments, found wood formation in *Pinus strobus* L. was correlated with Csupply. Stem regions with presumed increased C-supply experienced increased radial growth, mainly due to higher numbers of cells, with fewer cells produced in regions with reduced supply. Despite large changes in growth, labile carbohydrate concentrations were relatively stable across the different stem regions, reflecting homeostasis, with respiration rates correlated with C-supply. Rademacher et al. (2022) used chilling to control phloem transport in mature *Acer rubrum* L., and also found large effects on radial growth. The phloem chilling treatment also resulted in increased leaf carbohydrate concentrations and in downregulation of photosynthetic capacity during the peak growing season (Rademacher et al. 2022).

Despite overwhelming evidence that plants operate as integrated wholes, with source and sink activities feeding back on one another to maintain sugar homeostasis, global vegetation dynamics models (known as ‘Dynamic Global Vegetation Models’: DGVMs), which are used as components of Earth system models (Arora et al. 2020), and stand-alone as tools to analyse the global carbon budget (Sitch et al. 2024), assume a uni-directional relationship (sometimes with buffering from a storage pool). Growth is therefore treated as the direct outcome, and directly proportional to, the supply of C from photosynthesis, net of mitochondrial respiration (Friend et al. 2019). Conceptually this can result in unrealistic model behaviour for three broad, interrelated, reasons: (1) rates of source and sink activities in real plants respond differently to environmental factors, with sinks more sensitive than sources; (2) growth rate at any one time is ultimately limited by sink capacity, determined by meristem size and potential rates of cell proliferation and wall synthesis, which are under metabolic controls; and (3) if growth does not use all available C substrate, concentrations of labile C will increase and lead to down-regulation of source activities. However, the implications of C source-sink feedbacks, their dependencies, and how they can be implemented in DGVMs, remain unclear.

Various frameworks have been suggested for integrating C sources and sinks together within plant growth models (e.g. Friend et al. 2019; Jones et al. 2020; Gessler and Zweifel 2024), but these have not been formally assessed. Hayat et al. (2017) reviewed tree growth models with explicit consideration of sink processes (e.g. Schiestl-Aalto et al. 2015), and while there had been many advances, an approach integrating sources and sinks into a single framework was lacking. They therefore proposed and tested a new approach in which tree growth occurs from apical and lateral meristems, consuming C derived from photosynthesis, with a dynamic labile storage pool. The emphasis of the approach was on the geometric properties of the tree as it grows (i.e. height versus diameter), and controls thereof. It did not consider homeostasis of labile C or feedbacks between sources and sinks through controls on activities, although it did consider limitation to growth by C supply.

Other more recent models with explicit treatment of source and sink activities include that of Oswald and Aubrey (2023). Their approach is conceptually similar to the one described here, with its emphasis on feedbacks regulating carbohydrate concentrations and implementation using a parsimonious mathematical approach. However, it uses different formulations and focuses on the seasonal dynamics of starch concentrations in response to supply and demand, rather than the effects of sugar feedbacks on growth responses to environmental factors in DGVMs as here.

Formulating plausible approaches for the incorporation of C source-sink feedbacks within DGVMs and assessing potential consequences is therefore a priority. A parsimonious theoretical approach to a biological process or system can test and further understanding, as well as provide a context for more complete and mechanistic treatments. This paper takes such an approach to explore the implications of non-linear C source-sink coupling, through a common sugar pool, for C assimilation and growth responses to potential source and sink activities. ‘Potential’ here refers to the activity of the source or sink process when fully activated by the C-feedback, and is determined by the volume of the organ/tissue, its intrinsic metabolic machinery, and the effects of environmental and other non-C-feedback factors (e.g. phytohormones). We use the term ‘capacity’ to refer to the theoretical activity of the source or sink when not limited by C-feedback or any other signalling or environmental factors. The capacity is therefore determined only by the size of the source or sink tissues and their intrinsic metabolic machinery. This conceptual distinction between capacity and potential activity is formalised mathematically below. The primary aim of the work described here is to present and evaluate a plausible approach for incorporating source-sink interactions in a tree growth model at a conceptual level, rather than provide specific predictions. Indeed, the approach described here omits many processes that would be necessary to make detailed comparisons with specific experimental studies. Rather, we are interested in the qualitative behaviour of the approach in relation to the plausibilty of the model assumptions, and very much agree with John Thornley that ‘*[w]hile the predictions of a model must be examined and found acceptable, the most meaningful validation of a mechanistic model is at the level of its assumptions*’ (Thornley 2002). Nevertheless, we do compare simulations with and without C source-sink feedbacks with a set of elevated CO_2_ experiments in order to assess the general plausibility of our approach. We address the following two broad questions:

1) How can C source-sink feedbacks be represented in a tree growth model?

2) How does the emergent behaviour of a homeostatic approach to simulating tree growth compare with a traditional C-source-only-driven approach?

Below we describe the approach to be considered, determine a realistic set of parameter values, and explore time-dependent and equilibrium behaviours. Finally, we discuss the implications of considering C source-sink feedbacks in DGVMs in the manner proposed, make some suggestions for experiments to further investigate the model’s assumptions and implications, and describe the next steps that will be required to move beyond C-source-only-driven DGVMs.

## 2 Description

### 2.1 Governing equations

To gain insight into the consequences of C source-sink feedbacks for plant net photosynthesis and growth responses to environmental factors, and to explore a potential approach for their inclusion in DGVMs, it is assumed that C sources and sinks are linked by a common, single, sugar pool which can vary in size over time. The sugar is assumed to be sucrose, the source net photosynthesis, and the sink growth of structural mass (e.g. wood). The concentration of plant sucrose evolves with time according to,

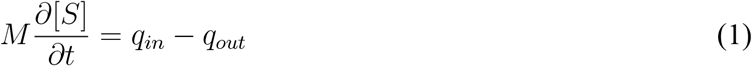

where [*S*] is the whole-plant sucrose concentration over all structural mass (g[Su] g[DM]^*−*1^; ‘Su’ = sucrose and ‘DM’ = dry matter), *q*_*in*_ is the source term (kg[sucrose] tree^*−*1^ day^*−*1^), *q*_*out*_ the sink term (kg[sucrose] tree^*−*1^ day^*−*1^), and *M* is total structural dry mass (kg[DM] tree^*−*1^).

Homeostasis requires coordinated adjustments to C source and sink rates in response to [*S*]. These adjustments are assumed to be achieved by sucrose inhibiting net photosynthesis and activating structural growth, as represented in Figure 1.

**Figure 1:**
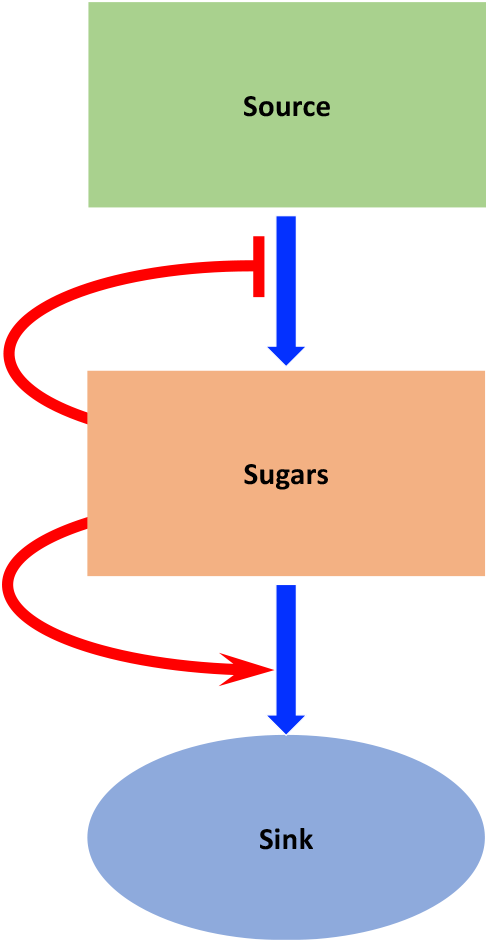
Schematic of whole-plant C source-sink feedbacks mediated by sugars. Blue arrows represent flows of sugars, the red flat-headed arrow represents an inhibition signal, and the red pointed arrow represents an activation signal. Sugars inhibit source activity and promote sink activity, resulting in homeostasis in their own concentration.

The functional forms of the feedbacks represented by the red arrows in Figure 1 are approximated by Hill functions (Hill 1910; Gesztelyi et al. 2012). Hill functions are frequently used to represent biologicalcontrol interactions (e.g. transcription factor regulation of gene expression: Alon 2020). They are biologically interpretable as a dose-response function where ‘cooperativity’ plays an important role. Cooperativity refers to the unit number of regulator required to initiate a unit molecular response, and is a frequent property of gene transcription regulation (Alon 2020). Sucrose is known to act as both a transcriptional and a translational regulator (Lastdrager et al. 2014), and affects the phospho-proteome through the protein kinase complex SnRK1 (Leene et al. 2022). Because these control pathways ultimately result in upor down-regulation of coordinated transcription factor networks, Hill functions are likely to be an appropriate way to represent source-sink interactions, albeit at a highly-aggregated level. Evaluation of different approaches showed that the continuous and sigmoidal form of the Hill functions as used here provides greater stability than, for example, linear functions, and have more biological realism, yet only have two parameters.

The Hill function describing the activating effect of sucrose on sinks, *f*_*a*_ (scalar), is given by,

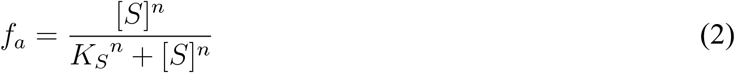

where *K*_*S*_ is the value of [*S*] at which sink activity is half its potential (i.e. fully activated) rate and the exponent *n* is the ‘Hill coefficient’, which determines the strength of the control (and is related to the cooperativity of the interaction, with higher values producing stronger control).

For simplicity, spatial variation in sucrose concentrations and finite transport between organs and tissues is not considered. However, it is acknowledged that spatial variation in concentrations (Furze et al. 2019) and transport limitations (Aluko et al. 2021), especially in large plants such as mature trees, will influence source-sink relations, and indeed play key roles in some models of C partitioning between organs such as that of Thornley (1972). However, a mean sucrose concentration is assumed to be an adequate indicator of whole-plant source-sink balance for our purposes, where we are interested in the general behaviour of the system under a set of broad assumptions, using a balanced depth of treatment across processes.

The inhibiting effect of sucrose on sources, *f*_*i*_ (scalar), is given by,

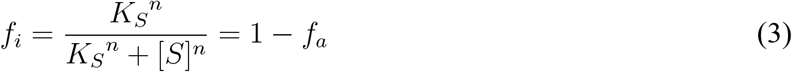

We assume linear effects of these feedback strengths on source and sink activities (i.e. the realised rates under a given set of conditions). Source activity, *q*_*in*_, is computed from the the sucrose feedback, any environmental constraints, and source capacity, which is assumed to linearly scale with the cross-sectional sapwood area at breast height,

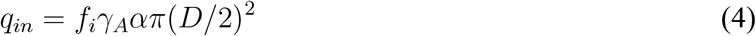

where *γ*_*A*_ is the environmental modulation of source activity (scalar), *α* is the source capacity per unit sapwood area (kg[sucrose] cm[sapwood]^*−*2^ day^*−*1^), and *D* is stem diameter at breast height (cm). For simplicity, respiration is not considered explicitly. Source activity is therefore equated with net photosynthesis (this is equivalent to assuming respiration is a fixed fraction of gross photosynthesis, although we recognise the limitations of this assumption: Thornley (2011)). Also, to maintain simplicity, heartwood is not considered.

*α* is the rate of source activity per unit sapwood area when there is no modulation by sucrose feedbacks or environmental factors. This capacity is assumed to be a constant in the analysis presented here, remaining unchanged over time and with tree size and/or age.

In order to remain parsimonious, sink activity is limited to consideration of growth in above-ground structure (below-ground mass likely scales relatively linearly with that above-ground, and therefore this assumption does not affect the general behaviour of the model), which is related to diameter at breast height allometrically,

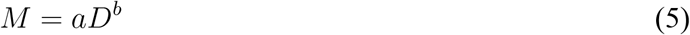

where *M* is total structural mass (kg[DM] tree^*−*1^) and *a* and *b* are empirical constants. Sink activity, *q*_*out*_, the change in structural mass over time (litter production is not considered), is therefore related to the change in *D* over time according to,

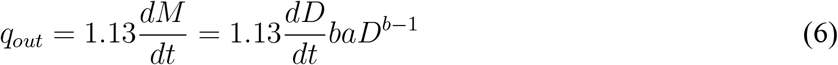

The additional constant converts DM to sucrose mass (i.e. 1.13 g[Su] g[DM]^*−*1^ = 0.474 g[C] g[DM]^*−*1^ / 0.421 g[C] g[S]^*−*1^, with the C content of DM taken from measurements in Panamanian rainforest tree species (Martin and Thomas (2011)).

The rate of increase in *D* with time is modulated by environmental factors and the sucrose activation factor,

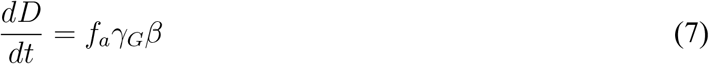

where *γ*_*G*_ is the environmental modulation of diameter growth (scalar) and *β* is the diameter growth capacity (cm day^*−*1^). As for source capacity (*α*), the sink capacity, captured by *β*, is assumed constant. The direct environmental factors, captured by *γ*_*G*_, include variables such as temperature and hydraulic status.

From Eqns. 2 and 3 it is evident that *K*_*S*_ can be considered as the neutral sucrose concentration in that the activities of the sources and sinks are equally activated when [*S*] = *K*_*S*_. At this neutral sucrose concentration, the sources and sinks are neither inhibited nor activated relative to each other, proceeding at their default rates. Neutral values of *f*_*i*_ and *f*_*a*_ are both 0.5 (scalar), and therefore *α* and *β* will need to be set to 2x their neutral rates under optimal environmental conditions (i.e. their rates when sugars are at the neutral value, *K*_*S*_, and the environmental controls, *γ*_*A*_ and *γ*_*G*_, = 1). If *f*_*i*_ and *f*_*a*_ were fixed at their neutral values, source and sink activities would be able to vary independently from one another in response to direct environmental factors resulting in non-homeostatic variation in sucrose concentrations. While not explicitly treated here, in real plants the effect of *f*_*i*_ on source activity (*q*_*in*_) could be realised through, for example, controls on the quantity of photosynthetic apparatus per unit foliage area (including stomatal density) and its activity through expression of photosynthetic genes (e.g. Sheen 1990), sucrose-induced stomatal control (Kottapalli et al. 2018) and/or controls on foliage area itself (Paul and Foyer 2001). Similarly, the effect of *f*_*a*_ on diameter growth could be realised through regulation of cambial cell proliferation, volume increase, and/or wall deposition (e.g. Kiba et al. 2019; Avidan et al. 2024).

*f*_*i*_ and *f*_*a*_ are acclimative controls that influence the sucrose balance through regulation of the activity of the sources and sinks. These effects are ultimately bounded by other factors, internal and external, captured by *α, β, γ*_*A*_, and *γ*_*G*_. We use various implementations of the model in order to investigate its behaviour. As well as simulations using the full (‘default’) model, in which structural mass changes significantly with time, we also analyse model behaviour with fixed mass (and hence fixed source and sink capacities) and equilibrium sucrose, resulting in additional insights. We also examine simulations with feedbacks switched off and in which growth is directly proportional to net photosynthesis as in traditional DGVMs. We refer to implementations with fixed mass and equilibrium sucrose as equilibrium simulations.

### 2.2 CO_2_ response

We investigate the influence of C source-sink feedbacks by simulating the growth of a tree germinating in 1700 CE, to match the mean age of the trees when the measurements were made used for model calibration (see below). Source activity (net photosynthesis) is assumed to vary in response to changing atmospheric CO_2_ over time, captured in the *γ*_*A*_ variable. Annual-mean historical atmospheric CO_2_ mixing ratio is taken from the ‘TRENDY’ forcings (Sitch et al. 2024). Values from 2000 CE to 2100 CE are taken from the SSP 585 scenario, reaching 1135.21 ppm in 2100 CE. This high-end scenario is used in order to examine a wide range of changed source activity.

The dependence of source activity on atmospheric CO_2_ is assumed to conform to a Michaelis-Mententype relationship,

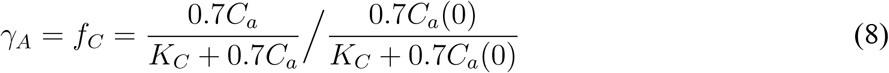

where *f*_*C*_ is the effect of CO_2_ on net photosynthesis (scalar), *C*_*a*_ is atmospheric CO_2_ mixing ratio (ppm), *C*_*a*_(0) is *C*_*a*_ a baseline value of atmospheric CO_2_ (assumed to be 368.9 ppm, the value in 2000 CE), and *K*_*C*_ is a constant. The constant 0.7 accounts for the reduction in CO_2_ between the atmosphere and internal air spaces of the leaves. When other environmental constraints are not considered the resulting value of *γ*_*A*_ in 2100 CE is 1.702.

The value of *K*_*C*_ is calibrated assuming that net photosynthesis responds to CO_2_ similarly to the response of net primary productivity (NPP). Friend (2010) modelled an increase in mean broadleaf evergreen tree NPP of 45% when CO_2_ increased from 375.7 ppm to 720 ppm, which is obtained if *K*_*C*_ = 400 ppm in Eqn. 8. The resulting response to CO_2_ using this value is shown in the SI, Figure 8. Norby et al. (2005) reported a mean stimulation of NPP of 23% across FACE experiments when CO_2_ was increased from 376 ppm to 550 ppm. Eqn. 8 gives a stimulation of 23.6% for the same change in *C*_*a*_.

### 2.3 Non-CO_2_ parameter values

The model is implemented here with a set of parameter values to simulate the growth of a single tree in a tropical evergreen rainforest climate without interference from neighbours. This minimises potential environmental constraints and avoids having to consider changing light availability with size. However, the general conclusions should apply to other situations, environments, and tree types. The set of default parameter values is given in Table 1.

**Table 1:** Parameters and their default values. ‘DM’ refers so structural dry matter and ‘Su’ refers to sucrose.

| Description | Symbol | Value | Units |
| --- | --- | --- | --- |
| $M(D)$ allometry coefficient (Eqn. 5) | $a$ | 0.1502 | kg[DM] cm<sup>-1</sup> |
| $M(D)$ allometry coefficient (Eqn. 5) | $b$ | 2.476 | unitless |
| CO<sub>2</sub> response constant (Eqn. 8) | $K_C$ | 400 | ppm |
| Neutral sucrose concentration (Eqns. 2, 3) | $K_S$ | 0.0288 | g[Su] g[DM]<sup>-1</sup> |
| Hill coefficient (Eqns. 2, 3) | $n$ | 4 | unitless |
| Source (net photosynthesis) capacity (Eqn. 4) | $\alpha$ | 0.0178/365 | kg[Su] cm[sapwood]<sup>-2</sup> day<sup>-1</sup> |
| Sink (growth) capacity (Eqn. 7) | $\beta$ | 0.658/365 | cm day<sup>-1</sup> |
| Timestep | $\Delta t$ | 1 | day |

#### 2.3.1 Parameter values derived from the literature

The allometric scaling of *D* to *M* (Eqn. 5) is derived from the pantropical relationshp between aboveground biomass (AGB) and diameter and height (*H*) of Chave et al. (2015). Height is calculated from *D* using the formulation of Feldpausch et al. (2011) for pantropical trees, which gives *H* = 30 m for *D* = 60 cm. Vieira et al. (2005) determined mean *D* growth rate at a tropical rainforest site ca. 90 km north of Manaus, Brazil, of ca. 0.2 cm yr^*−*1^and a median tree age of 300 yr, and therefore *D* = 60 cm is typical for a mature tree at this site. Vieira et al. (2005) describe the forest canopy height as ca. 35 m and as relatively homogeneous. This height-diameter relationshiop is similar to the relationship in Feldpausch et al. (2011). Therefore the Feldpausch et al. (2011) formulation was used to replace *H* in the AGB equation of Chave et al. (2015). Wood density is used in the *M* (*D*) relationship of Chave et al. (2015) and is assumed here to equal 0.670 g[DM] cm^*−*3^ (Nogueira et al. 2005; mean value at the site near Manaus). Combining these assumptions, and simplifying to Eqn. 5, yields *a* = 0.1502 kg[DM] cm^*−*1^ and *b* = 2.476. For *D* = 60 cm this gives *M* = 3796 kg[DM].

The value for *K*_*S*_ (Eqns. 2, 3) is taken as 2.88% of structural dry matter, the mean sucrose concentration combining foliage, branch, stem, coarse roots, and fine roots measured by Würth et al. (2005) across nine tropical forest species in Panama (individual compartment values were 4.0%, 3.9%, 2.6%, 2.3%, and 1.6% respectively). The Hill coefficient (*n*; Eqns. 2, 3) is set to 4, the top of the range given as typical for regulation of gene transcription by Alon (2020), producing a fairly steep function (i.e. relatively tight homeostatic regulation).

#### 2.3.2 Parameter values derived by calibration

The parameters *α* and *β* were calibrated to give the observed mean diameter growth at the rainforest site close to Manaus (i.e. *D* = 60 cm after 300 yr of growth: Vieira et al. 2005, assumed to be in 2000 CE), and the observed sucrose concentration in Panama (see above). The calibrated values are then *α* = 0.0178/365 kg[Su] cm[sapwood]^*−*2^ day^*−*1^ and *β* = 0.658/365 cm day^*−*1^. This calibration relies on the assumption that sucrose concentration is at the neutral value (*K*_*S*_) in 2000 CE, the assumed year of the observations (the implications of this assumption are assessed below).

### 2.4 Numerical method and initial values

The evolutions of the structural (Eqn. 6) and sucrose (Eqn. 1) masses over time were solved simultaneously using the classic Runge–Kutta method, with a 1 day timestep. The initial value for the stem diameter, *D*, was taken from Figure 1D of Melo et al. (2015), showing recently germinated *Genipa americana* L. seedlings, a wide-ranging tree species of Central and Southern America. The value obtained was 0.1 cm. The initial sucrose concentration was set to *K*_*S*_.

## 3 Results

### 3.1 Dynamic simulations with the default model

The results of a dynamic simulation with initial conditions as described above, the default set of parameter values (Table 1), CO_2_ 1700-2100 CE as described above, and with sucrose feedbacks activated and structural mass increasing over time, are shown by the solid blue lines in Figure 2.

**Figure 2:**
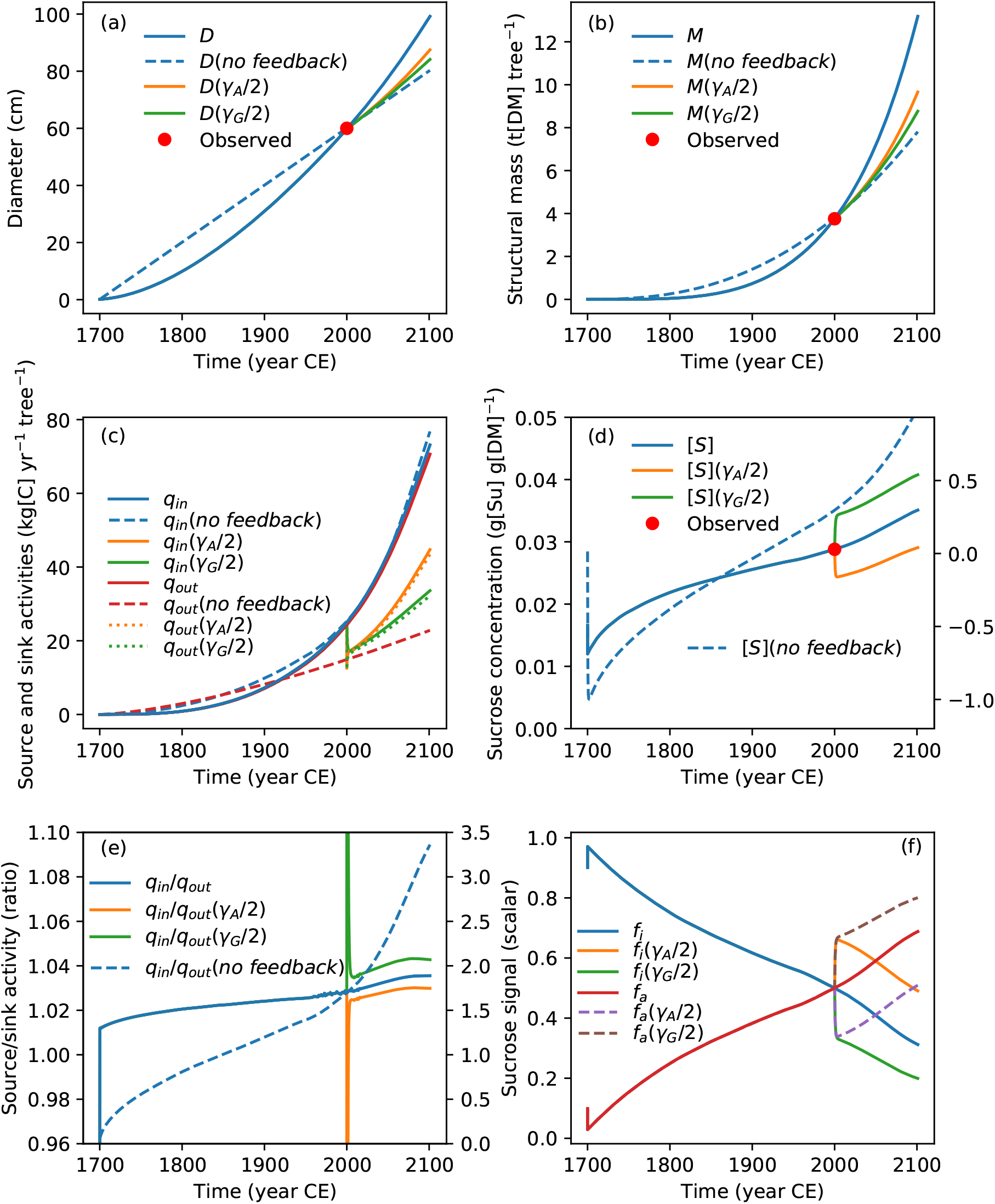
Simulated tree growth, source/sink activities, and sucrose dynamics using the default model (solid blue and red lines), when feedbacks are switched off by setting [*S*] = *K*_*S*_ (‘*nof eedback*’), and when source (‘*γ*_*A*_/2’) or sink (‘*γ*_*G*_/2’) activity is reduced by 50% from 2000 CE (simulating environmental constraints). (a) Diameter at breast height; (b) structural mass; (c) source (*q*_*in*_ = net photosynthesis) and sink (*q*_*out*_ = structural growth) activities; (d) sucrose concentration (N.B. the no-feedback case uses the right-hand y-axis); (e) ratio of source to sink activity (N.B. the no-feedback case uses the right-hand y-axis); and (f) sucrose signals. Observations as described in main text; in (d) positioned for both y-axes.

Diameter at breast height increases exponentially, passing through the observed calibration value in 2000 CE (Fig. 2a). Structural mass increases strongly with time due to the allometric relationship relating *M* to *D* in Eqn. 5 (Fig. 2b; solid blue line; cf. Stephenson et al. (2014)).

Source and sink activities increase in parallel with each other, but the rate of net photosynthesis is always slightly higher than the rate of growth (Fig. 2c; solid red and blue lines). After an initial decline, the sucrose concentration increases rapidly for the first ca. 100 yr, from a value of 0.012 g[Su] g[DM]^*−*1^, subsequently saturating as it approaches *K*_*S*_, before increasing more steeply again from around 1956 CE, reaching 0.0351 g[Su] g[DM]^*−*1^ in 2100 CE, 21.9% above the 2000 CE calibration neutral value of 0.0288 g[Su] g[DM]^*−*1^ (Fig. 2d). The ratio of source to sink activities remains above 1 after an initial rapid adjustment, and increasing slightly throughout the simulation (Fig. 2e), driving the increase in sucrose concentration. Increasing sucrose concentration causes declining source and increasing sink activation control signals, with neutrality occurring in 2000 CE as calibrated (Fig. 2f; solid blue and red lines).

#### 3.1.1 No-feedback simulation

The dynamics of the full default simulation, shown by the solid blue and red lines in Fig. 2, are driven by a combination of the changing allometry of the tree as it grows, increasing CO_2_ mixing ratio, and feedbacks on source and sink activities through sucrose concentrations. The marginal effects of the sucrose feedbacks were determined by repeating the full default simulation but with sucrose feedbacks switched off, achieved by setting [*S*] = *K*_*S*_ in Eqns. 2 and 3. Actual sucrose mass was still allowed to vary as a function of inputs and outputs (Eqn. 1; negative values were allowed), but not affect source or sink activity. In order to facilitate comparison with the default simulation, *β* was recalibrated to 0.4/365 cm day^*−*1^, forcing *D* = 60 cm in 2000 CE.. The resulting behaviour is shown by the dashed lines in Fig. 2a-e.

Without sucrose feedbacks, diameter increases linearly at the mean observed rate (Fig. 2a; dashed blue line). Therefore the exponential increase in *D* with time in the default simulation must be due to the increasing concentration of sucrose promoting sink activity (solid blue line in Fig. 2d). Growth in structural mass is still exponential, due to the allometric scaling with *D* (Eqn. 5), but with a substantially smaller exponent, resulting in 40.1% lower mass by the end of the simulation than when sources and sinks feed back on one another (Fig. 2b; dashed blue line).

Source activity follows a similar trajectory to that in the default simulation, although with slightly higher values (Fig. 2c; dashed blue line). In contrast, the rate of growth, *q*_*out*_, is strongly affected by the sucrose feedback (Fig. 2c; red solid and dashed lines). When [*S*] = *K*_*S*_, growth rate increases almost linearly due to the linear increase in *D*, falling behind that of the default simulation during 1920 CE. This reinforces the interpretation that, in the default model, increasing sucrose concentrations drive the strong exponential increase in diameter (and hence mass) growth rates.

Fig. 2d shows the trajectory of sucrose concentration for the no-feedback simulation (right-hand y-axis, positioned such that the observation symbol is correct for both y-axes; blue dashed line). Values are negative prior to 1912 CE due to growth demands exceeding source supply when the tree is small (Fig. 2c; dashed lines). This demonstrates the success of the sucrose feedbacks in simultaneously controlling concentrations homeostatically while allowing realistic growth. When feedbacks are not activated, realistic growth results in impossible sucrose concentrations.

The increase in the ratio of source to sink activity with time is much more pronounced when feedbacks are switched off than in the default model (Fig. 2e; right-hand y-axis; dashed blue line). The overall trend with time is due to both the changing ratio between source and sink capacities with tree size and increasing CO_2_. The acceleration in the rate of increase after ca. 1950 CE is due to the lack of compensating feedback on source activity, increasing sensitivity to CO_2_. The relevance of changing source and sink capacities was confirmed by a strong increase in the source/sink ratio even when CO_2_ is fixed at the value in 1700 CE, and with feedbacks switched off (not shown). While source capacity scales as the square of diameter (Eqn. 4), sink capacity scales with diameter to the power of ca. 1.5 (i.e. *b −* 1; Eqn. 6). Fixing CO_2_ and increasing *b* to 3 eliminates the trend in source/sink ratio (not shown). Therefore, growth becomes less source-limited with size for purely allometric reasons, which, in the full default model, are incompletely compensated by inhibition of source activity and activation of sink activity as the tree grows (Fig. 2f; solid red and blue lines).

#### 3.1.2 Effect of CO_2_ on simulated dynamics using the default model

The consequences of increasing CO_2_ for default model behaviour were investigated. Simulation results are shown in (Fig. 3) for increasing (solid lines) and fixed (dashed lines) CO_2_.

**Figure 3:**
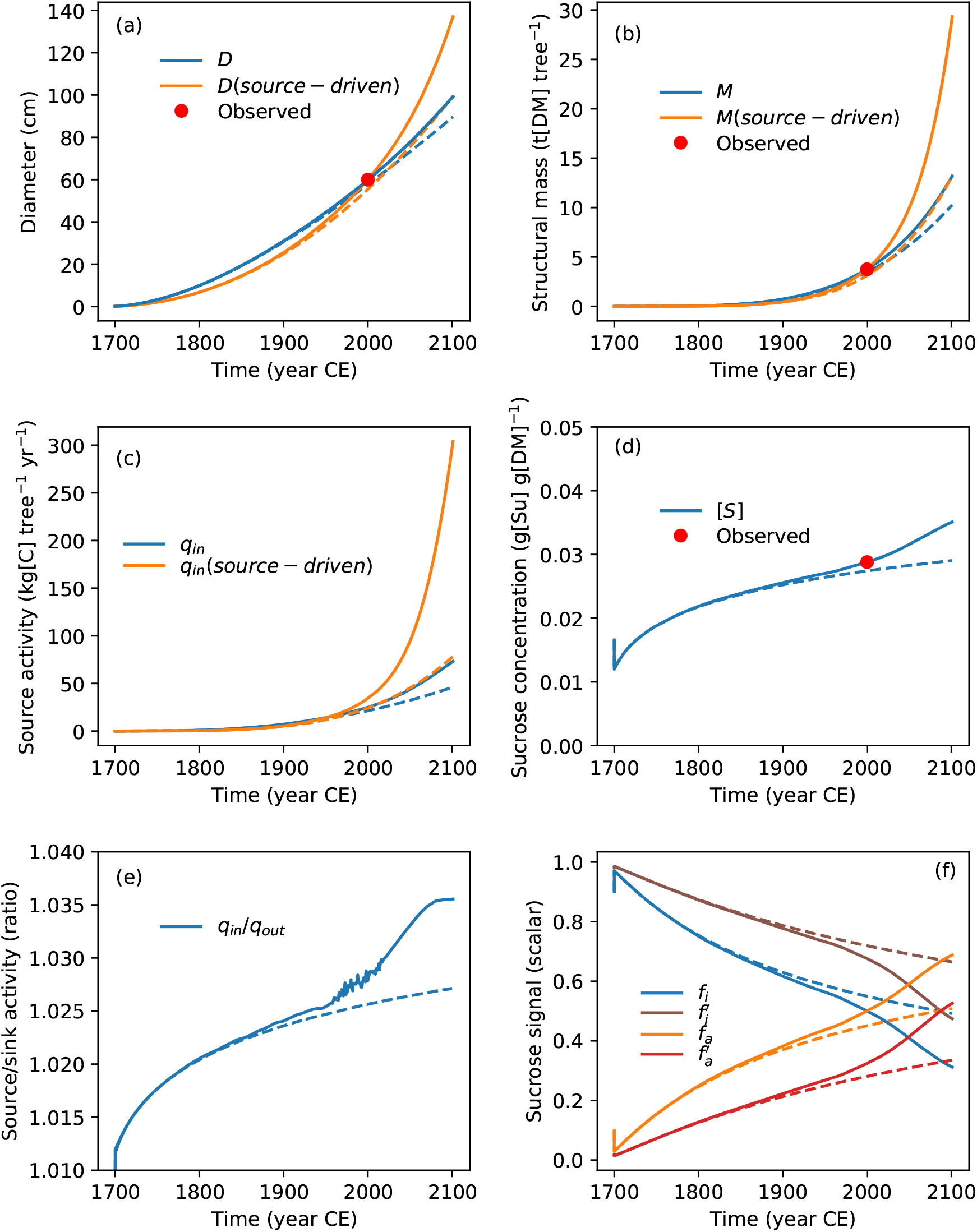
Simulated tree growth, source activity, and sucrose dynamics with (solid lines) and without (dashed lines) increasing atmospheric CO_2_. Results are shown for the default model, when source-driven (i.e. *q*_*out*_ = *q*_*in*_, ‘*source − driven*’), and when *K*_*S*_ is increased by 20% (symbols with apostrophes in (f)). (a) Diameter at breast height; (b) structural mass; (c) source activity per tree; (d) sucrose concentration; (e) ratio of source to sink activity; and (f) sucrose signals. Observed values as in Fig. 2.

Fixing CO_2_ at its mixing ratio in 1700 CE (i.e. 277.75 ppm) throughout the simulation shows that the growth stimulation due to CO_2_ in the default simulation causes *D* at the end of 2000 CE to be increased slightly, from 58.0 cm to 60.1 cm, and *M* by 9.3% (Fig. 3a,b; solid and dashed blue lines) The stimulation of *M* increases to 28.5% by the end of 2100 CE (Fig. 3b).

Increasing CO_2_ causes both source and sink activities to be 17.4% higher by the end of 2000 CE, and 58.5% higher by the end of 2100 CE (Fig. 3c; sink and source activities are indistinguishable at the resolution of the figure, hence only source is shown). Increasing CO_2_ causes sucrose concentration to be 20.9% higher (i.e. an increase from 0.0290 g[Su] g[DM]^*−*1^ to 0.0351 g[Su] g[DM]^*−*1^) at the end of 2100 CE (Fig. 3d), indicating incomplete buffering. When CO_2_ is fixed, sucrose concentration approaches an asymptote, and the ratio of source to sink activity also saturates (Fig. 3e). Higher sucrose concentrations with increasing CO_2_ lead to greater source inhibition and sink activation than when CO_2_ is fixed. With fixed CO_2_, neutrality is delayed from 2000 CE until 2084 CE (Fig. 3f).

#### 3.1.3 Consequences of source-sink coupling for responses to CO_2_

The consequences of source-sink coupling for modelled responses to CO_2_ were investigated by comparing the behaviour of the default model with that of a version using assumptions similar to traditional C sourceonly-driven models. Two simulations were performed, both with and without increasing CO_2_, in which growth was assumed to equal net photosynthesis. This was achieved by setting *q*_*out*_ = *q*_*in*_ and [*S*] = *K*_*S*_. *γ*_*A*_ was calibrated to 1.36x*f*_*C*_ in order for *D* to reach 60 cm in 2000 CE for increasing CO_2_, facilitating comparison with the default simulation.

Increasing CO_2_ enhances source and sink activities by 41.5% at the end of 2000 CE when feedbacks are not considered, compared to an enhancement of only 17.4% when they are (Fig. 3c; orange and blue lines), an increase in sensitivity of 142.4%. This change in sensitivity increases to ca.5x by the end of 2100 CE. In other words, source-sink feedbacks reduce the response of net photosynthesis and of growth rate to increasing CO_2_ by 58.7% by the end of 2000 CE, and by 80.2% by the end of 2100 CE.

Structural mass is affected similarly. By the end of 2000 CE, the growth stimulation due to CO_2_ for the no-feedback simulation increases *M* by 22.4%, increasing to 121.9% by the end of 2100 CE (Fig. 3b). The response of mass growth to projected CO_2_ to the end of the century is therefore 5.4x greater (i.e. 121.9/22.4) when growth is directly source-driven compared to when source and sink are coupled through sucrose feedbacks. In other words, source-sink feedbacks reduced the simulated mass increase due to increasing CO_2_ by the end of the century by 76.7%.

In the C source-only-driven simulation, the lack of feedback inhibition on net photosynthesis leads to greater *D* growth, which increases the source capacity (Eqn. 4), increasing *D* further. *D* is 137.0 cm at the end of the source-only-driven simulation when CO_2_ increases, compared to 99.3 cm without increasing CO_2_ (Fig. 3a). The resulting difference in sapwood area increases source capacity by 90.4%. In comparison, at the end of the default simulations, increasing CO_2_ enhances source capacity by only 22.4%, which together with feedback inhibition of-37.6% from neutrality (i.e. *f*_*i*_ = 0.312) (Fig. 3f) accounts for the much lower CO_2_ response in the default model.

These comparisons are quantified for responses at the whole-tree level, which will tend to be greater than those on a per-unit ground area basis due to the relationships between source activity and size through Eqn. 4, and sink activity and size through Eqn. 6. Normalising source and sink activity by sapwood area removes this effect of size, and the resulting trends in source activity are shown in Fig. 4.

**Figure 4:**
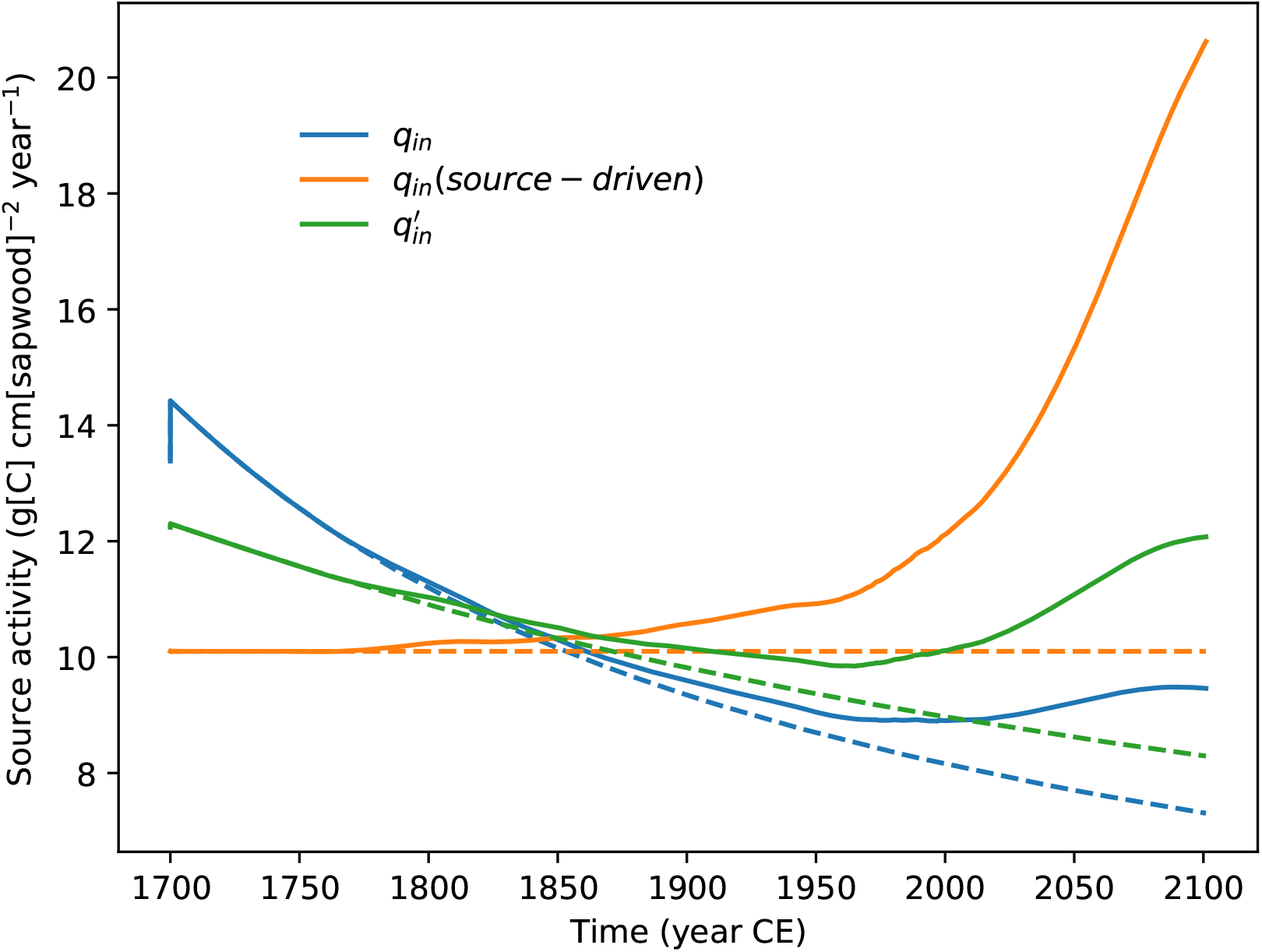
Simulated source activity per unit sapwood area with (solid lines) and without (dashed lines) increasing atmospheric CO_2_. Results are shown for the default model, when source-driven (i.e. *q*_*out*_ = *q*_*in*_, ‘*source − driven*’), and when *K*_*S*_ is increased by 20% (symbol with apostrophe).

Sapwood-area-normalised source activity declines strongly as the tree grows using the default model until the latter half of the 20th Century, when increasing CO_2_ halts this trend (Fig. 4; solid blue line). In contrast, the sapwood-normalised source rate increases with time when growth equals net photosynthesis, especially in the final 100 yr or so with increasing CO_2_ (Fig. 4; solid orange line). The decline in the sapwood-normalised rate with time in the default simulation is therefore due to feedback inhibition of net photosynthesis (Fig. 3f; solid blue line).

Increasing CO_2_ in the default simulation increases the sapwood area-normalised rate of net photosynthesis by 29.5% by the end of the simulation compared to when CO_2_ does not increase (Fig. 4; solid and dashed blue lines). The comparable effect for the source-only-driven simulation is to increase source activity by 104.2% (Fig. 4; solid and dashed orange lines), which is the change in the CO_2_ forcing, *f*_*C*_ (Eqn. 8). In other words, the response of the sapwood-area-normalised rate of source activity by the end of 2100 CE to increasing CO_2_ is reduced by 71.7% due to the source-sink feedbacks. This reduction is similar to the non-sapwood-area-normalised reduction of 79.7%, showing that the majority of the feedback effect on the response is direct rather than due to changing allometry with size.

#### 3.1.4 Growth responses to perturbed source versus sink activities

The model separates environmental controls on source and sink activities through *γ*_*A*_ and *γ*_*G*_, respectively. The transient response of the default model to environmental controls was evaluated by decreasing either *γ*_*A*_ or *γ*_*G*_ by 50% at the end of 2000 CE, and maintaining the perturbed value to the end of 2100 CE. The resulting behaviours are compared with the unperturbed simulation in Fig. 2.

*D* and *M* growth are both reduced for both perturbations, as expected, but reducing *γ*_*G*_, the environmental control on sink activity, has a greater effect (Fig. 2a,b; orange and green solid lines). Reducing source activity reduces sucrose concentration (Fig. 2d; solid orange line), which promotes source activity and inhibits sink activity compared to the default simulation (Fig. 2f; solid orange and dashed purple lines). Conversely, reducing sink activity leads to an increase in sucrose concentration (Fig. 2d; solid green line), inhibiting source and promoting sink activity (Fig. 2f; solid green and dashed brown lines). An asymmetry in these responses is evident in Figure 2d, which is due to the forms of the Hill functions, as discussed below in relation to starvation versus growth constraints. The 50% reduction in source activity due to environmental constraints results in a 38.8% reduction in growth rate by the end of 2100 CE, whereas a 50% reduction in sink activity due to environmental constraints results in a 54.1% reduction in final growth rate (i.e. 33.1% greater effect (Fig. 2c)).

#### 3.1.5 Sensitivity to *K*_*S*_

The simulations described above used the somewhat arbitrary assumptions that, in 2000 CE, *K*_*S*_ = 0.0288 g[Su] g[DM]^*−*1^ and sources and sinks are in balance. There is no *a priori* reason why these assumptions need to be the case, and indeed the change in [*S*] over the 400 yr of simulated growth (Fig. 2d) suggests other values of *K*_*S*_ would be compatible with [*S*] = 0.0288 g[Su] g[DM]^*−*1^ in 2000 CE. As the aim of the work described here is not to determine precise predictions (and indeed omits many processes that might be expected to affect tree growth, such as competition), but rather to investigate the relative implications of considering C source-sink feedbacks, the pertinent question to ask is therefore to what extent are the conclusions affected by the chosen value of *K*_*S*_? A secondary question then is how can we determine *K*_*S*_, or is there even a single value over the lifetime of a tree?

To address the first question, two further simulations, with and without increasing CO_2_, were performed. These were similar to the default simulations, except that the value of *K*_*S*_ was increased by 20% to 0.0336 g[Su] g[DM]^*−*1^), and the *α* and *β* parameters re-calibrated to give *D* = 60 cm and [*S*] = 0.0288 g[Su] g[DM]^*−*1^ in 2000 CE. Overall growth to 2100 CE is increased, with *M* 24.8% higher and the rate of growth 52.6% higher. Sucrose concentrations are little changed, and therefore the greater overall growth is due to higher values of *f*_*i*_ throughout the simulation, shown in Fig. 3f (solid brown line), with [*S*] remaining below *K*_*S*_ until late in the simulation. These higher values of *f*_*i*_ drive significantly higher source and hence sink activities, while their ratio is not changed.

Comparing simulations with CO_2_ fixed at its value in 1700 with those driven by increasing CO_2_, but both with 1.2x*K*_*S*_, gives a CO_2_-driven increase in final *M* of 47.6%, compared to 28.5% using the default value of *K*_*S*_. The equivalent comparison for the final rate of net photosynthesis is 99.4% vs. 58.5%. Hence the value of *K*_*S*_ has a large impact on predicted growth response to CO_2_. Higher values of *K*_*S*_ increase the CO_2_ response because the tree has a greater requirement for C in order to reach the neutral value, which is not achieved until nearly 2100 CE for increasing CO_2_ when *K*_*S*_ = 0.0336 g[Su] g[DM]^*−*1^, shown by the crossing of the solid brown and red lines in 3f).

### 3.2 Equilibrium condition

#### 3.2.1 Insights from analytical solution

The model equilibrium condition, with fixed structural mass and equilibrium sucrose, allows insights into model sensitivity to parameters and impacts of source-sink feedbacks in the absence of confounding effects such as changing allometry. It is also amenable to analytical solutions, allowing complete understanding. Non-equilibrium states are harder to evaluate as behaviour depends on distance from equilibrium. We define the equilibrium state as one where source and sink activities are in balance, and therefore,

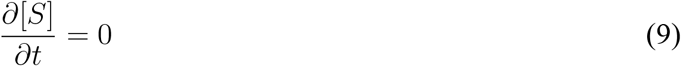

and mass is fixed. Fixing mass removes the confounding effect of changes in source and sink capacities with size, and dilution of sucrose as mass increases with growth. Taking into account sucrose feedbacks (Equations 2 and 3), the response of equilibrium net photosynthesis and growth rate to potential source and sink activities (and by implication also capacities) reduces to (see Appendix 9.2 for derivation),

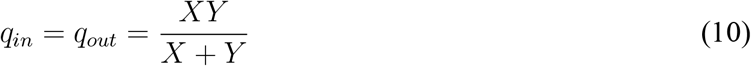

where *X* is the potential source activity, equal to the ratio of *q*_*in*_ to *f*_*i*_, and *Y* is the potential sink activity, equal to the ratio of *q*_*out*_ to *f*_*a*_. These potential activities are those of the source or sink when at full activation (i.e. *f*_*i*_ or *f*_*a*_ = 1) (N.B. activation refers to the sucrose signal effect, not the influence of environmental factors, which might cause the potential rate to be lower than the capacity). Eqn. 10 shows that, when sucrose concentration and mass are at equilibrium, the response of growth rate to potential growth or net photosynthesis rates is independent of the strength of the feedbacks (i.e. the Hill coefficient, *n*) and the value of the neutral (equilibrium) sucrose concentration (i.e. *K*_*S*_), and equal whether potential source or sink activities (and capacities) are changed. It also shows that the responses of the realised source and sinks rates to changes in either potential source or sink activity are equal.

The change in growth rate with change in potential source activity reduces to (see Appendix 9.2 for derivation),

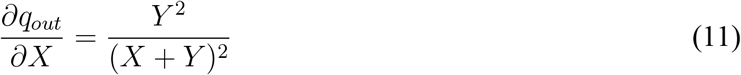

indicating a steep increase in growth rate with potential source activity at low source capacity, gradually saturating as potential source activity increases. If potential source and sink activities are equal, then 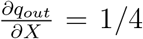, indicating a strongly constrained response of growth to potential source activity (and therefore source capacity) for the equilibrium state.

#### 3.2.2 Responses of rates and state to potential source and sink activities

The responses of the equilbrium model rates and state to variation in potential source or sink activities are shown in Figure 5 using the set of default parameters in Table 1, with *D* = 60 cm (and therefore *M* = 3796 kg[DM]). Potential source or sink activities were varied using a scalar applied to *q*_*in*_ (Eqn. 4) or *q*_*out*_ (Eqn. 6), respectively.

**Figure 5:**
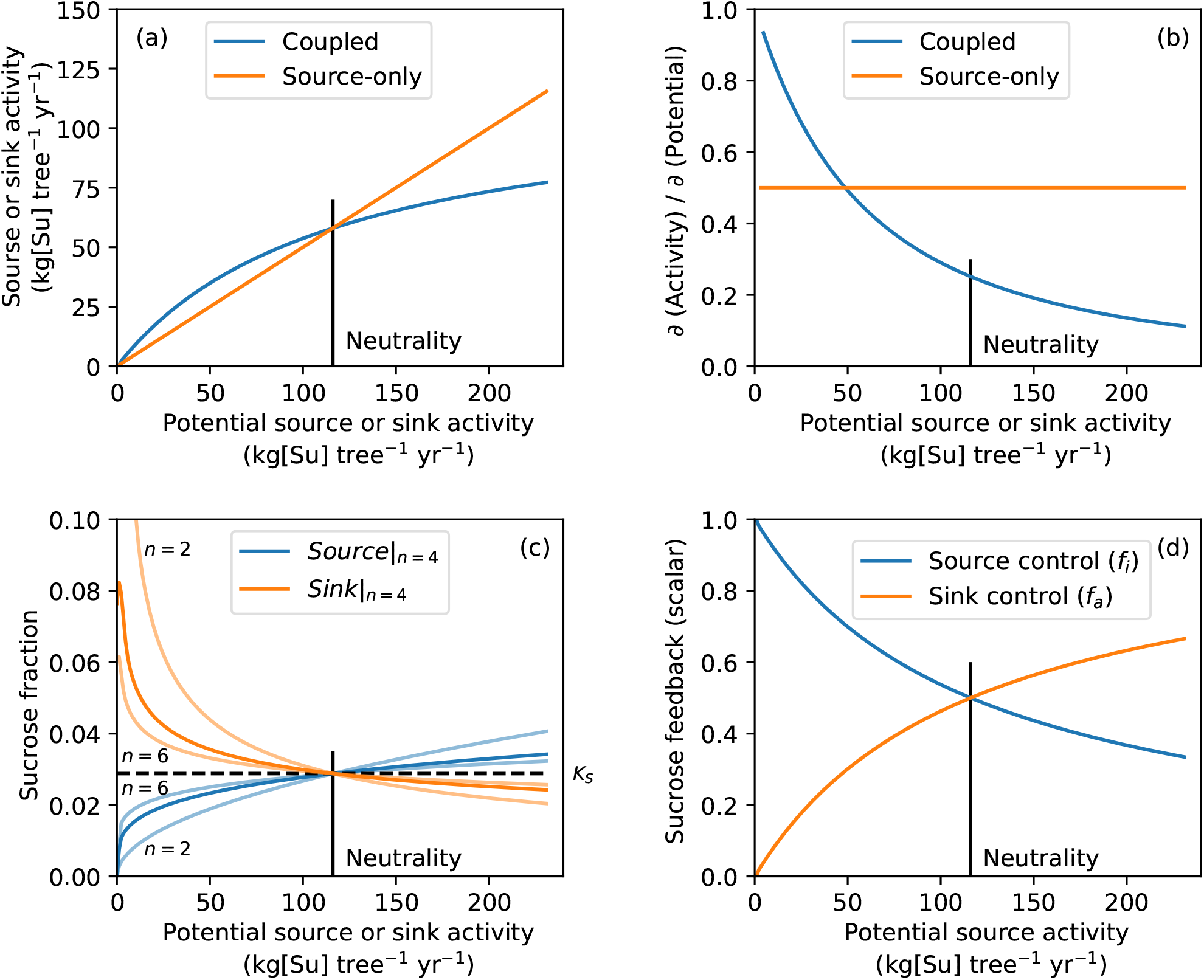
Simulated equilibrium model rates and state using parameter values in Table 1, except where specified, with *D* = 60 cm. (a) Response of equilibrium source and sink activities to potential source and sink activities (‘Coupled’). Also shown is the response assuming sink activity is equal to source activity with no feedback effects (‘Source-only’). In this case the ‘potential’ activities on the y-axis are simply double the actual activities in order to force the response through the same value as the coupled implementation under the neutral condition. (b) Derivatives of the responses shown in (a). (c) Responses of equilibrium sucrose concentration to potential source and sink activities using different values of Hill coefficient (*n*). Also shown is the value of *K*_*S*_, horizontal dashed line. (d) Responses of equilibrium sucrose feedback controls to potential source activity. All panels indicate the position of the neutral condition for the coupled model (i.e. when *f*_*a*_ = *f*_*i*_ = 0.5 and [*S*] = *K*_*S*_) using a vertical black line.

The responses of equilibrium source (net photosynthesis) and sink (growth) rates to varying potential source or sink activities are shown in Figure 5a. By definition the two rates are equal at equilibrium. Also, the responses are the same whether potential source or sink activities are varied (cf Eqn. 10, and therefore only one curve is shown (‘Coupled’). This curve shows strong saturation with increasing potential source or sink activity, demonstrating the constraining effect of sources on sinks and sinks on sources, each imposing an upper limit on the other (cf. Equation 10). Even when potential source or sink activity is 2x its default value, growth or net photosynthesis, respectively, increase to only 1/3 above their default rates, 2/3 of their potential rates. Equilibrium growth has the upper limit of *Y* as *X* is increased (obviously the growth rate cannot exceed its potential), and vice versa (Equation 10).

Also shown is the response of the growth rate if it is assumed to equal net photosynthesis with no feedbacks (‘Source-only’), similar to assumptions in C-source-centric approaches as for the non-equilibrium simulations. This response is plotted assuming that the source activity is 50% of its potential in order to facilitate comparison with the coupled model. This causes the source-only simulation to equal the coupled model at the neutral condition (i.e. when *f*_*a*_ = *f*_*i*_ = 0.5 and [*S*] = *K*_*S*_ in the coupled model), which requires net photosynthesis to be 1/2 of its potential rate. Above this neutral condition, the two approaches diverge considerably as the effect of source capacity on growth saturates due to the limiting rate of potential growth in the coupled model. At double the default potential source activity, the growth rate is 50% higher without feedbacks than with homeostatic coupling, and therefore the coupling reduces growth by 1/3 compared to the non-coupled assumption.

The derivatives of the responses in Figure 5a are shown in Figure 5b. At the neutral point for the coupled model, the sensitivity of growth to variation in potential source activity is only 50% of the sensitivity if growth is entirely source-driven. The sensitivity of the coupled model only reaches that of the source-driven approach when potential source activity is reduced to 50% of its default value.

Equilibrium sucrose concentration increases with potential source activity and decreases with potential sink activity, equalling *K*_*S*_ at their default values (Figure 5c; dark blue and orange lines). Source-sink feedbacks are highly effective at constraining sucrose levels over a wide range of potential source and sink rates. For example, a 50% increase in potential source activity from its default value results in [*S*] increasing by only ca.10%. Similarly, an increase in potential growth activity of 50% reduces [*S*] by only ca.7%. In contrast, as potential source or sink activities approach zero, sucrose concentration deviates considerably from the equilibrium value.

Figure 5c also shows the equilibrium sucrose responses for different values of the Hill coefficient (*n*). Lower values are less effective at constraining sucrose levels, but the outcome is little changed between *n* = 4 and *n* = 6. *n* could be calibrated from observed behaviour in sucrose concentration with respect to environmental conditions such as atmospheric CO_2_.

Figure 5d shows the responses of the feedback signals to potential source activity for the equilibrium model. Controls are stronger when potential source activity is below its default value than above, conforming to a starvation response. The responses to changing potential sink activity are the inverse of these responses, with source control increasing, and sink control decreasing, with sink strength (not shown).

#### 3.2.3 Flux responses to starvation or halted growth with fixed structural mass

The dynamic consequences for the model, with fixed structural mass as in the previous section, of halted source or sink activities were investigated. These scenarios represent either constrained net photosynthesis (‘Starvation’; e.g. due to defoliation) or constrained growth (‘No growth’; e.g. due to intense drought directly affecting xylogenesis, e.g. Eckes-Shephard et al. (2021)), respectively. For both, the default model was run using the parameters in Table 1 for four model years to bring it to equilibrium, starting from the initial values given above and with *D* fixed at 60 cm. For the starvation scenario, the C source was then switched off by setting *q*_*in*_ = 0. The resulting behaviour of the growth sink (*q*_*out*_) and sucrose pool ([*S*]) are shown in Figure 6.

**Figure 6:**
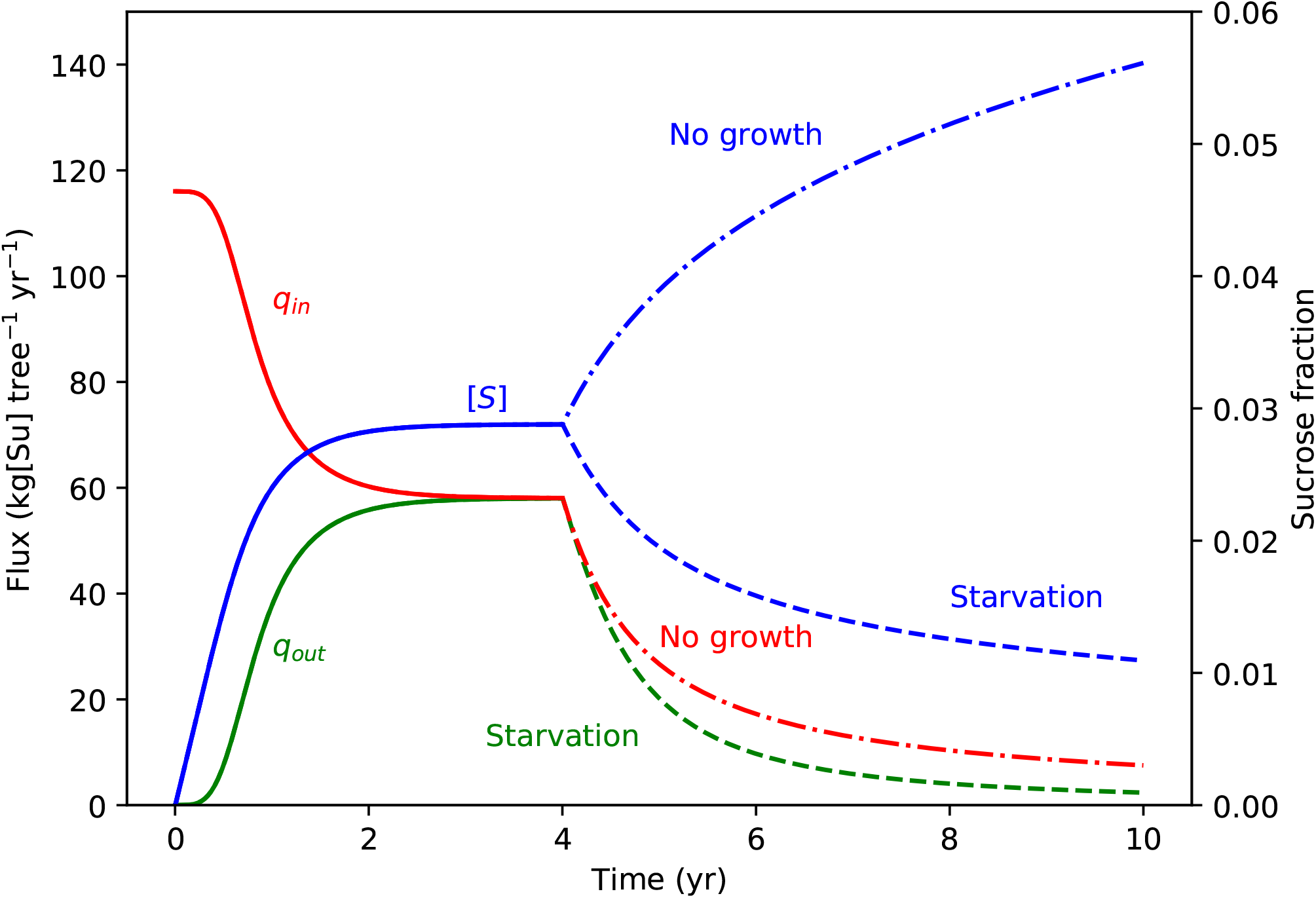
Simulated dynamic source and sink fluxes and sucrose concentrations using parameter values in Table 1 and with fixed structural mass. A spin-up of four years is followed by either *q*_*in*_ (‘Starvation’) or *q*_*out*_ (‘No growth’) set to 0.

Growth declines as sucrose concentrations fall, taking 312 days to reach 50% of the equilibrium rate, *>* 10 months. This long timescale results from the large initial mass of sucrose and the sucrose feedback reducing sink strength as sucrose declines. Such a response is consistent with the observed effects of defoliation on radial increment in mature trees, with severely defoliated trees exhibiting 40-60% reduction in stem growth during the year of defoliation (e.g. Naidoo and Lechowicz 2001; Hilmers et al. 2023). The continuation of declining sucrose but sustained growth with starvation is also consistent with observations (e.g. Wang et al. 2021). The long-term gradual decline in growth rate is a consequence of the forms of the Hill functions. Different formulations could be used if Hill functions do not capture observed temporal dynamics, but at present such a test is limited by the availability of relevant experimental results.

For testing the effect of constrained growth, the C sink was switched off by setting *q*_*out*_ = 0 at the end of the spin-up period. The effect of this imposed growth restriction is to cause a reduction in net photosynthesis (*q*_*in*_), with a somewhat slower dynamic than the response of growth to zero net photosynthesis (Figure 6). This asymmetry in effects of source and sink constraints is due to inherent asymmetry within the Hill functions. The low limits are reached when sucrose declines to zero, but a numerically equivalent absolute increase from the neutral value will result in rates still somewhat below their maxima. For example, halving [*S*] from its neutral value results in *f*_*a*_ falling by 88.2%, whereas increasing [*S*] by the same absolute amount causes *f*_*i*_ to fall by 67.0%. Hence declining sucrose levels have a stronger relative effect than increasing levels, resulting in the asymmetry between the effects of starvation and no growth in Figure 6. Even at the end of the simulation period, after six years of zero growth, net photosynthesis is 13.0% of its neutral value, whereas after the same period with no net photosynthesis, growth has declined to only 4.1% of its neutral value.

Observational evidence for effects of restricted growth independently of external effects on photosynthesis is rare due to methodological challenges. Using phloem chilling in mature *Acer rubrum*, Rademacher et al. (2022) found reduced woody growth below the chilling collars, accumulation of sugars in phloem and leaves above the collars, and reduced photosynthesis capacity, all consistent with the behaviour of the model under *q*_*out*_ = 0. The changed sucrose concentrations found in these experiments also occur in the model.

### 3.3 Comparison to free-air CO_2_ enrichment (FACE) experiments

Direct comparison between the model output and observations is challenging because the model simulates the growth of the above-ground woody material of an individual tree free from interactions with neighbours, and without consideration of numerous other processes and conditions relevant to specific experimental responses. Figure 7 shows observed increases in total tree growth rates (NPP) across a range of FACE experiments, together with model outputs, plotted against tree age. Model outputs are shown both per tree and per unit sapwood area, whereas all observed responses are per unit ground area and with closed canopies. Despite the lack of site-level calibration, the simulated growth-rate enhancements per unit sapwood area capture both the absolute and relative observed enhancements per ground area across tree age for the majority of the observations (green and red lines). The exception is AspenFACE, with a growth enhancement closer to that predicted at the whole-tree level (blue and orange lines). The treatment plots in this experiment were exposed to increased CO_2_ from tree germination, and the results reported from canopy closure, 1-3 yr later. Despite canopy closure, the majority of the growth enhancement could be explained by increased light interception (Norby et al. 2005), and therefore it seems reasonable that the growth response should be closer to that predicted at the whole-tree level.

**Figure 7:**
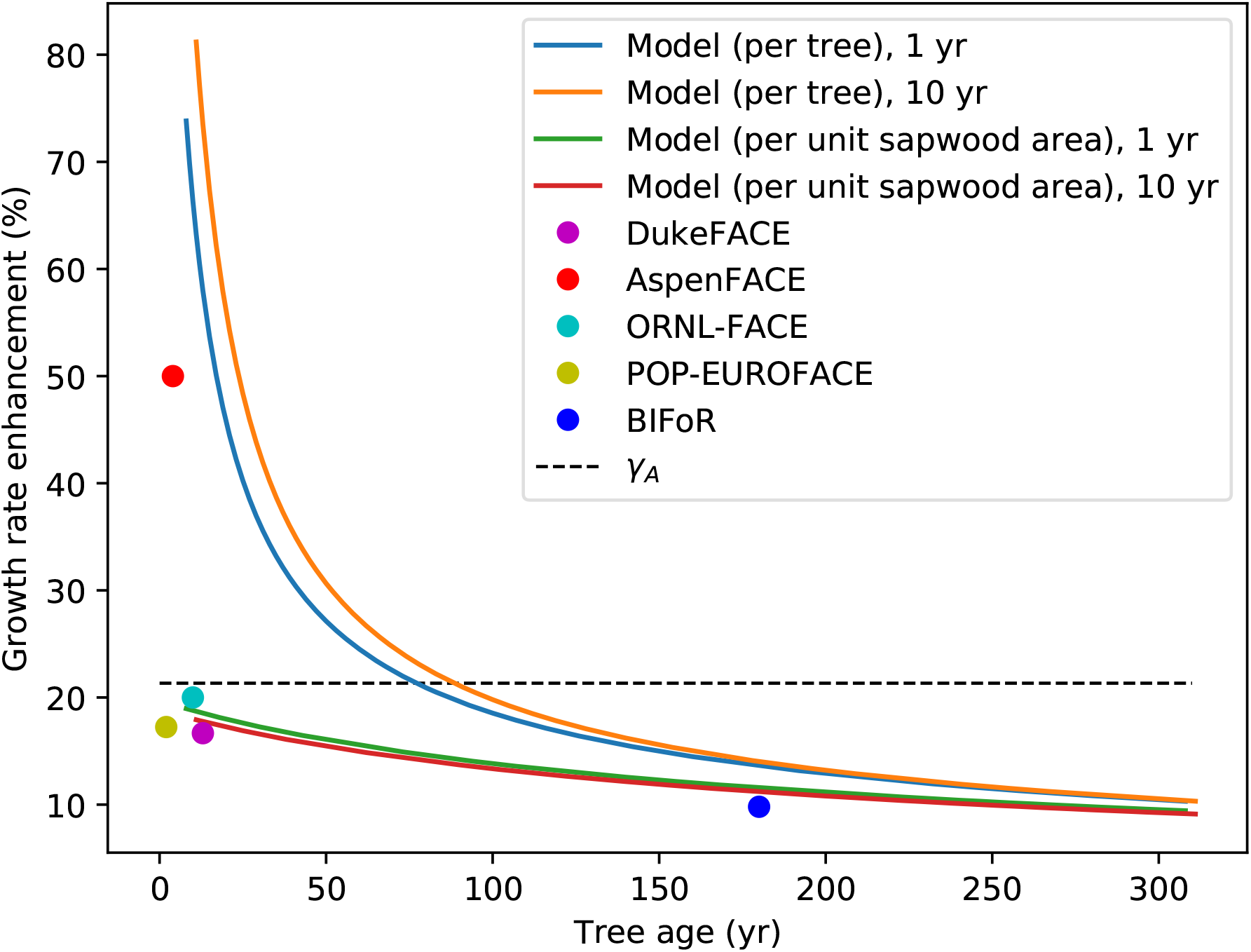
Simulated and observed growth-rate enhancement due to elevated CO_2_. Model responses derived using full default model across a range of germination dates, with or without +165 ppm CO_2_ added to the natural CO_2_ concentration from 2000 CE. Growth enhancement is the percentage increase in growth rate due to the additional CO_2_ in either 2001 CE (‘1 yr’) or 2010 CE (‘10 yr’). BiFOR observations taken from Norby et al. (2024). Other observations extracted from Figure 1A of Norby et al. (2005) and related to ages derived from Table 1 and main text therein. *γ*_*A*_ is the increase in potential source rate per unit sapwood area due to added CO_2_ in 2000 CE (Eqn 8).

**Figure 8:**
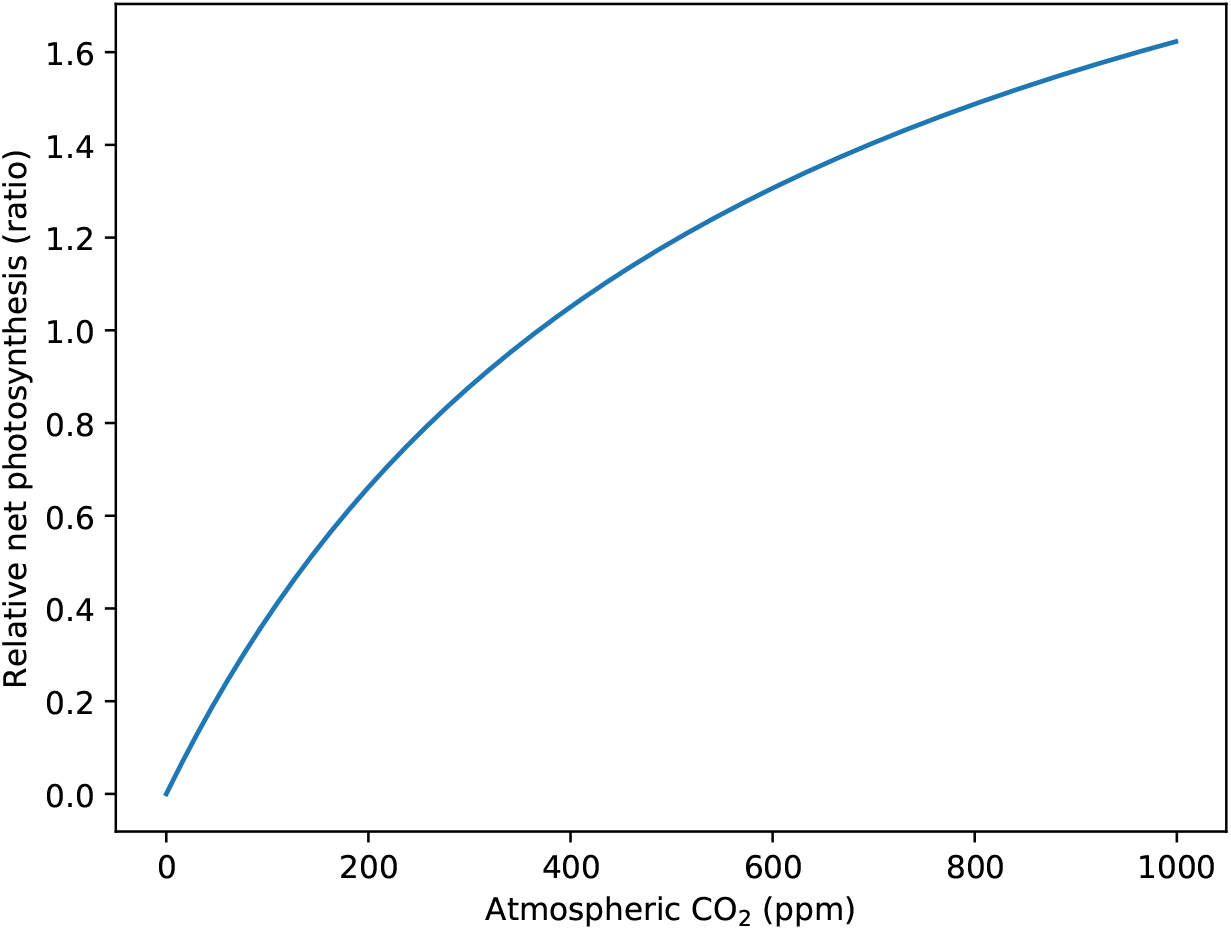
Calibrated response of net photosynthesis to atmospheric CO_2_ as described by Eqn. 8. Response normalised to 1 at *C*_*a*_ = 368.87 ppm, the mean observed value in 2000 CE.

The predicted growth enhancement declines with tree age due to changing allometry, as discussed above. Greater potential source activity per tree with size results in greater inhibition of net photosynthesis through sucrose feedback. The growth enhancement per unit sapwood area approaches *γ*_*A*_ as age approaches 0, declining with age to approximately the same degree as observed (Fig. 7).

## 4 Discussion

Despite the knowledge that plants operate as integrated wholes, with strong coupling between sources and sinks, plant growth in DGVMs is traditionally treated as being a direct, one-way, function of C-source activity. While there have been numerous calls for the inclusion of explicit growth processes in DGVMs (e.g. Fatichi et al. 2019; Friend et al. 2019), approaches both for doing this, and for coupling growth to source activity, require development and assessment. Here an approach based on the concept of homeostasis is described and analysed, with feedbacks from a common sucrose pool regulating source and sink activities through Hill functions.

Using a set of parameters appropriate for a tree growing in a tropical rainforest environment, the approach described here is effective at maintaining sucrose homeostasis while allowing for independent environmental effects on source and sink rates and increasing source and sink capacities as the tree grows. Sucrose feedbacks scale potential (i.e. fully activated) source and sink rates under given environmental conditions, allowing the effects of external environmental factors and size (which affects whole-tree source and sink capacities) to be treated while the system is still able to behave homeostatically. Maintenance of homeostasis through feedbacks constrains the extent to which changes in potential source or sink rates can affect overall growth (cf. Figure 5a). Notably, the effect of increasing net photosynthesis on growth is strongly constrained by the potential growth rate, leading to feedback-inhibition of source activity.

An important implication of these results is that the strong CO_2_-fertilization effects on growth predicted by many DGVMs (e.g. Huntzinger et al. 2017) are unlikely to be realistic, with major consequences for projections of the future terrestrial C sink and ecosystem functioning. Compared with the traditional assumption of C-source-only-driven growth, the response of the rate of growth to rising atmospheric CO_2_ projected for the end of this century is reduced by ca.70% (i.e. from +104% to +29% for the sapwoodarea-normalised rates) due to the effects of feedbacks. The response of biomass in 2100 CE to increasing CO_2_ is reduced by ca.77% (i.e. from +122% to +29%).

These findings are reflected in experimental results, with growth stimulation less than expected given the (initial) stimulation of photosynthesis due to, e.g., elevated CO_2_ (Körner 2006). The negative sucrose feedback on source activity reduces predicted growth enhancement, with greater reductions in larger trees, consistent with observations (cf. Figure 7). Variation in the growth response to CO_2_ between studies is often attributed to different levels of nutrient availability, which limits biomass production due to stoichiometric constraints (e.g. Jiang et al. 2024). This constraint is incorporated into a growing number of DGVMs (e.g. Zaehle and Friend 2010), but the consequences of dynamic coupling of sinks and sources has not been explored. Our approach allows for the incorporation of stoichiometric constraints through the *β* parameter, with feedback on source activity producing homeostasis in sucrose concentrations. This then allows for a treatment of the mechanistic interactions between C and nutrient constraints, such as of N and P, which is more physiologically realistic than current approaches (e.g. Zaehle and Friend (2010)). The responses of the model to C starvation or halted growth provide insights regarding the temporal dynamics of the sugar pool (Figure 6). It is noteworthy that these response times are of the same order as a typical growing season in mid-latitude temperate climates. This suggests that total labile carbohydrates in mature trees will be dynamically and constantly converging to their equilibrium value, getting close to equilibrium during the growing season regardless of the starting point. However, a full assessment of the dynamics of carbohydrates during the course of the growing season (and interactions with phenology) requires separation of sugars and starch, and compartments within trees such as branches and roots with transport between them (cf. Furze et al. 2019).

Inclusion of explicit growth in DGVMs has been advocated largely on the basis of evidence that growth processes can be more strongly regulated by environmental factors, such as soil moisture and temperatures away from optima, than photosynthesis (e.g. Gessler and Zweifel 2024). Coupling sources and sinks as described here broadens this justification through demonstrating that, even if sources are more strongly environmentally regulated than sinks, the growth response will still depend on feedbacks and hence sink capacity (which is never infinitely plastic), meaning growth processes need to be represented for realistic simulation of C uptake. Growth is not just a consequence of (net) photosynthesis, but is rather a function of the dynamics of the whole-plant non-structural carbohydrate pool, which is itself a function of growth and C-source rates.

The homeostatic behaviour of the sucrose pool underpinning our model is consistent with observations and theory (e.g. Rademacher et al. 2021; Miret et al. 2024). Therefore this component of the approach presented here is presumably at least qualitatively realistic. The specific details of the feedback functional forms are harder to verify, but the Hill coefficient does not affect the equilibrium responses to potential C source and sinks rates. However, the *K*_*S*_ parameter, the neutral sucrose concentration, has a large effect on model behaviour (cf. Figure 3), and the Hill coefficient affects the temporal dynamics under nonequilibrium conditions. It remains to be shown if other functional forms result in more realistic dynamic behaviour. An important next step is to determine the response functions relating source and sink activities to sucrose concentrations *in vivo*.

More generally, it would be valuable to explore experimentally the assumptions within, and insights derived from, the model presented here. In particular, the functional dynamics of homeostatic feedback, captured by Hill functions using *K*_*S*_ and *n* parameters, could be explored by manipulating source and sink strength and measuring carbohydrate concentrations over time (cf. Figure 6), while acknowledging the challenges this poses (Landhäusser et al. 2018). What are the response functions relating source and sink activities to sucrose concentrations? Do Hill functions reproduce the observed behaviour? It would be important to explore homeostatic behaviour over a range of plant sizes, with the expectation that larger plants would be more buffered and potentially relatively more sink-limited (cf. Figure 2; Hayat et al. 2017). Are *K*_*S*_ and *n* constant over size, time, and/or external conditions? How does *K*_*S*_ and *n* vary between species? What physiological mechanisms determine these functional forms? Another important consideration is the extent to which starch reserves buffer sucrose concentrations and whether starch synthesis is a passive overflow or actively controlled (cf. Hartmann and Trumbore 2016). Experimental manipulation of source and sink activities in the context of starch-sucrose dynamics would be extremely interesting. Starch-deficient mutant hybrid aspen trees are starting to be used for such experiments (Wang et al. 2022), and are likely to yield important information in future work.

The issue as to under which circumstances plants are C source-or sink-limited has been the subject of much discussion (e.g. Körner 2006). The analysis presented here, based on maintenance of sucrose homeostasis through feedbacks on sources and sinks, suggests that such a clear dichotomy does not reflect the dynamic and continuous nature of plant carbon physiology, where source and sink activities are dynamically regulated by both environmental and internal conditions, including the balance between source and sink demands through feedbacks. Growth and (net) photosynthesis are coupled. Their relative importance varies, however, as their potential rates vary, for example with size through allometric changes and/or with environmental conditions, which helps to explain why experimental evidence is inconclusive as to whether C source or sink limits growth (a similar sentiment was expressed by Gessler and Zweifel (2024)). In our model, the relative effect of source or sink strength can be quantified as the derivative of growth with respect to *α* or *β* (as in Figure 5(b)). C source is more important than sink when when *∂q*_*out*_/*∂α > ∂q*_*out*_/*∂β*, and vice versa, but neither is in complete control under either condition.

In conclusion, a parsimonious theoretical approach to coupling plant C source and sink activities, through a shared sucrose pool, has been presented and its implications assessed. Such coupling strongly constrains the response of growth to changes in C-source activity and the approach suggests a pathway to more realistic integration of whole-plant physiology and environmental controls in DGVMs. Future research will focus on extending this framework through developing the physiological mechanisms underlying source and sink activities (i.e. replacing *α* and *β* by process-based representations), treating whole-plant allocation, including below-ground and foliage dynamics, and including interactions between individuals through shading and shared soil water, leading to the integration of whole-plant coupled C-source-sink growth dynamics in a DGVM.

## 5 Acknowledgements

This work was supported by the Natural Environment Research Council through Grant NE/W000199/1. It was performed using resources provided by the Cambridge Service for Data Driven Discovery (CSD3) operated by the University of Cambridge Research Computing Service (www.csd3.cam.ac.uk), provided by Dell EMC and Intel using Tier-2 funding from the Engineering and Physical Sciences Research Council (capital grant EP/T022159/1), and DiRAC funding from the Science and Technology Facilities Council (www.dirac.ac.uk). PRT wishes to thank the Cambridge Mathematics Placements (CMP) programme, funded in part by donations made by corporate partners and individual donors to the CMP Supporters Club and by the Department of Geography, University of Cambridge. ADR and TR acknowledge support from the National Science Foundation, awards DEB-1741585 and DEB-1832210. PF acknowledges support from the Swiss National Science Foundation project CALEIDOSCOPE (212902). AHE-S acknowledges support from the European Research Council under the European Union Horizon 2020 Programme (grant no. 758873, TreeMort). This study contributes to the Strategic Research Areas BECC and MERGE.

## 7 Author contributions

ADF conceived of the study, wrote and ran the model, and wrote the first draft of the manuscript. PRT derived the stability analysis and wrote the relevant section in the Supporting Information, as well as contributing to mathematical ideas in the main manuscript. PF suggested the study on C starvation and halted growth. All authors contributed to interpretation of model output and provided input to the manuscript.

## 8 Data availability

Model code can be found on GitHub: https://github.com/adfriend45/sugar

## 9 Supporting Information

### 9.1 CO_2_ forcing

### 9.2 Equilibrium growth rate

Let potential source activity, *X*, be given by,

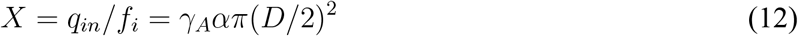

and potential sink activity, *Y*, be given by,

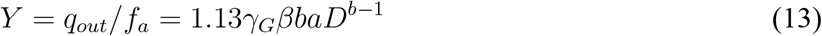

At equilibrium source and sink fluxes are equal,

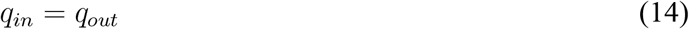

and therefore.

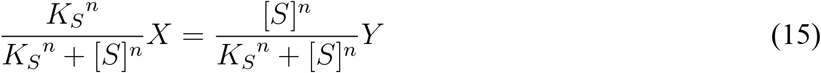

Solving for equilibrium [*S*]^*n*^ gives,

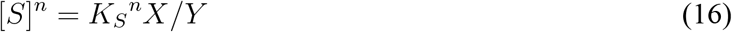

substituting into the expression for *q*_*out*_ and simplifying,

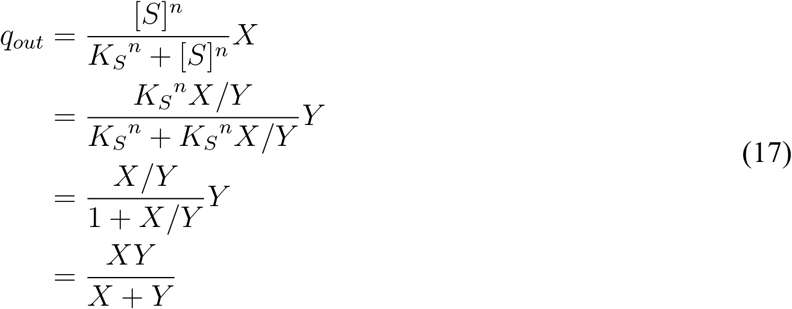

The derivative of *q*_*in*_ with respect to *X* can then be found using the quotient rule,

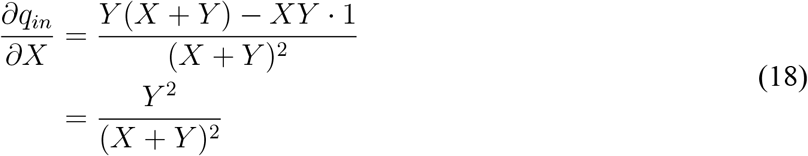

